# Optimizing Short-format Training: an International Consensus on Effective, Inclusive, and Career-spanning Professional Development in the Life Sciences and Beyond

**DOI:** 10.1101/2023.03.10.531570

**Authors:** Jason J. Williams, Rochelle E. Tractenberg, Bérénice Batut, Erin A. Becker, Anne M. Brown, Melissa L. Burke, Ben Busby, Nisha K. Cooch, Allissa A. Dillman, Samuel S. Donovan, Maria A. Doyle, Celia W.G. van Gelder, Christina R. Hall, Kate L. Hertweck, Kari L. Jordan, John R. Jungck, Ainsley R. Latour, Jessica M. Lindvall, Marta Lloret-Llinares, Gary S. McDowell, Rana Morris, Teresa Mourad, Amy Nisselle, Patricia Ordóñez, Lisanna Paladin, Patricia M. Palagi, Mahadeo A. Sukhai, Tracy K. Teal, Louise Woodley

**Author notes:** These authors contributed equally to this work **Emails:**. **Author Contributions:** J.J.W. authored the NSF 2026 Reinventing Scientific Talent proposal and with R.E.T. developed the conference proposal and secured funding. J.J.W. and R.E.T. contributed equally to the design and development of the conference and written outputs (this paper, www.bikeprinciples.org, implementation roadmap). B.B., S.S.D., K.J.L., T.M., T.K.T., C.vG. participated on the organizing committee and contributed to discussions and development of the participant nomination and scoring (J.J.W. & R.E.T. made final selections), as well as all meeting agendas. All other participants contributed to conference discussions, with A.L., G.S.M., L.P., M.S., and S.D. developing independent synthesis documents. Contributed conference presentations were delivered by M.B., S.D., C.R.H., K.L.H., J.J., A.L., J.L., K.L.J., R.M., T.M., P.O., L.P., P.P., M.S., T.K.T., R.E.T., C.vG., J.J.W., and Charla Lambert (see acknowledgements). N.C. compiled notes and provided suggestions for the development of the implementation roadmap. All authors contributed to the discussions and review of synthesized documents. All authors participated in review of the submitted manuscript.

## Abstract

Science, technology, engineering, mathematics, and medicine (STEMM) fields change rapidly and are increasingly interdisciplinary. Commonly, STEMM practitioners use short-format training (SFT) such as workshops and short courses for upskilling and reskilling, but unaddressed challenges limit SFT’s effectiveness and inclusiveness. Prior work, including the NSF 2026 Reinventing Scientific Talent proposal, called for addressing SFT challenges, and a diverse international group of experts in education, accessibility, and life sciences came together to do so. This paper describes the phenomenography and content analyses that produced a set of 14 actionable recommendations to systematically strengthen SFT. Recommendations were derived from findings in the educational sciences and the experiences of several of the largest life science SFT programs. Recommendations cover the breadth of SFT contexts and stakeholder groups and include actions for instructors (e.g., make equity and inclusion an ethical obligation), programs (e.g., centralize infrastructure for assessment and evaluation), as well as organizations and funders (e.g., professionalize training SFT instructors; deploy SFT to counter inequity). Recommendations are aligned into a purpose-built framework— “The Bicycle Principles”—that prioritizes evidenced-based teaching, inclusiveness, and equity, as well as the ability to scale, share, and sustain SFT. We also describe how the Bicycle Principles and recommendations are consistent with educational change theories and can overcome systemic barriers to delivering consistently effective, inclusive, and career-spanning SFT.

**SIGNIFICANCE STATEMENT:** STEMM practitioners need sustained and customized professional development to keep up with innovations. Short-format training (SFT) such as workshops and short-courses are relied upon widely but have unaddressed limitations. This project generated principles and recommendations to make SFT consistently effective, inclusive, and career-spanning. Optimizing SFT could broaden participation in STEMM by preparing practitioners more equitably with transformative skills. Better SFT would also serve members of the STEMM workforce who have several decades of productivity ahead, but who may not benefit from education reforms that predominantly focus on undergraduate STEMM. The Bicycle Principles and accompanying recommendations apply to any SFT instruction and may be especially useful in rapidly evolving and multidisciplinary fields such as artificial intelligence, genomics, and precision medicine.

## INTRODUCTION

A shared characteristic of science, technology, engineering, math, and medical (STEMM) disciplines is that “new technologies replace the skills and tasks originally learned by older graduates” and “technological progress erodes the value of these skills over time (1).”

For example, advanced computational methods such as machine learning have transformed life science with 1,487 publications on PubMed referencing this technique in 2012, compared to 30,684 in 2022 (2). This level of disruptive change can leave practitioners at risk of having large areas of their discipline rendered unintelligible to them (3–5). Life scientists see computational and data management training as their most unmet need (6, 7), reflecting the challenge in modern science to incorporate knowledge and skills from across multiple disciplines (e.g., computational methods, see (8)).

This project explored the application of evidence-based teaching and principles of inclusion and equity to improve short-format training (SFT) such as workshops, bootcamps, and short courses. SFT is widely used for upskilling and reskilling in rapidly evolving disciplines such as life science where disruptive changes and shifting skill sets are increasingly common. SFT’s popularity can be attributed to several positive features such as its relatively low cost and time commitment, as well as its capacity for rapid update and customization. Given the urgent need for full participation in STEMM (e.g., (9–11)), it is also important to note that SFT can be designed or revised to equitably include historically excluded people. For example, *The Carpentries Toolkit of IDEAS* provides strategies before, during, and after SFT to promote inclusion, diversity, equity, and accessibility (12). In addition to purely technical skills, SFT is also used to disseminate and reinforce professional practices such as research rigor, reproducibility, and other open science skills (13, 14).

Despite its positive features, SFT’s efficacy—its ability to measurably improve learners’ knowledge, skills, and abilities—may be much lower than is commonly realized. Feldon et al. (15) is the most extensive independent and peer-reviewed study to date that systematically evaluated the impact of SFT on life scientists. This study analyzed SFT interventions involving 294 life science Ph.D. students from 53 U.S. institutions across 115 variables and found “no evidence of effectiveness.” Feldon et al. concludes that “boot camps and other short formats may not durably impact student outcomes,” and that more effort and resources should be spent on improving SFT.

Feldon’s findings align with prior research in and beyond the U.S. (e.g., (16,17)). The 2022 *5^th^ Global Report on Adult Learning and Education* of the UNESCO Institute for Lifelong Learning (18) notes that only 60% of participating EU countries use learning outcomes as a quality measure of adult learning and education, across all types of instruction. Quality assessment is recognized to be difficult “…because of the diversity and plurality, and sometimes decentralized and deregulated nature, of the field—not to mention the variety of learners’ aims—across national and regional settings.” ((18) p. 25). We do not assert that all SFT is ineffective. However, we know from other STEMM instructional settings that some learners are likely to benefit from a learning opportunity no matter how well or how badly it is taught. Cooper et al. (19) notes that, “(a)lthough most STEM faculty and practicing scientists have learned successfully in a traditional format, they are the exception, not the norm, in their success” (p. 281). If instruction is only “effective” for learners who are unaffected by the quality of instruction, then that instruction is literally exclusionary because not all learners will benefit.

There is a strong rationale for reforming SFT. SFT’s positive features satisfy needs that are difficult or impossible to address otherwise. There is consistent demand for SFT training opportunities worldwide (e.g., (7)), and university, research institutes and government agencies continue to provide substantial funding for SFT; from 2017-2022 GrantExplorer reported expenditure of $4 billion in NSF, $83 million in NIH, and $767 million in DoD funding to projects associated with some SFT output (20–22).

Currently, STEMM education reform focuses primarily on formal higher education (FHE) (23–27), but these efforts are unlikely to directly impact SFT. Contrasting FHE and SFT (see Figure 1) and noting the variabilities identified by UNESCO (18) suggests that techniques used to improve FHE, if feasible for SFT, would likely require modification. SFT is not simply a “short” version of instruction in FHE; their only shared characteristic is that formal knowledge about teaching and learning applies (e.g., (28, 29)). Considering the features of FHE holistically, it should also be noted that FHE’s relative uniformity makes it easier to develop systemic reforms; the ability to address systemic problems is an additional reason FHE is more concretely improvable than SFT. As Reinholz et al. (30) suggests, “The goal of improving postsecondary STEM education requires careful attention to many interlocking systems and parts of systems.” (see also (31); p. 952; (32, 33)). Efforts to improve STEMM instruction in FHE have proceeded with some assurance that findings and interventions could be generalized across similar institutions and programs. This generality is more difficult for SFT where FHE’s “systems and parts of systems” may be difficult to compare, unrecognizable, or non-existent.

**Figure 1.**
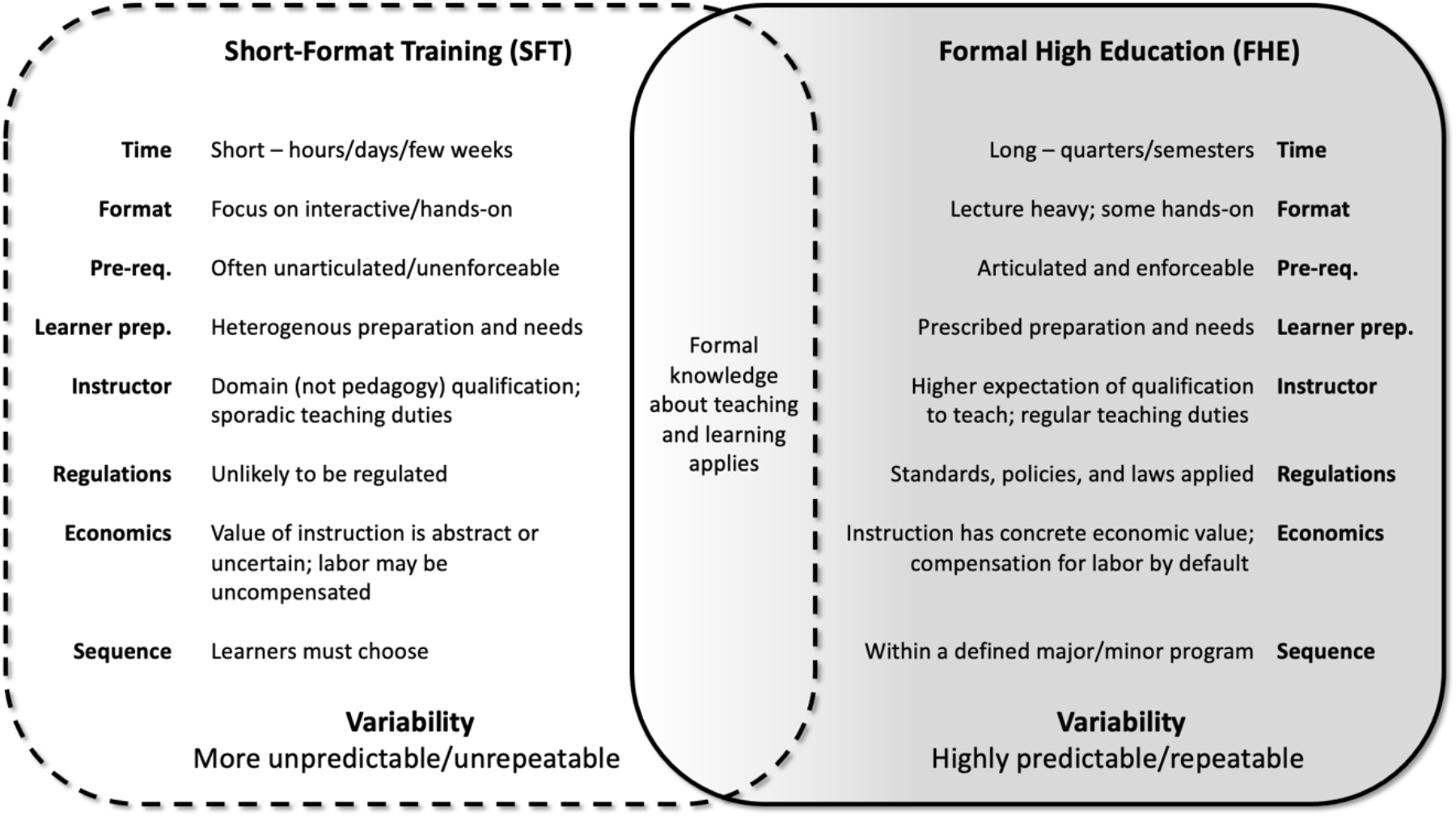
SFT is highly variable: contrasting features of short-format training (SFT) and formal higher education (FHE). We wrap SFT with a dashed-line highlighting that any given SFT may be difficult to define exactly (see expanded definition of SFT, Supplemental Information); FHE’s solid line indicates its greater uniformity and lower variation. The **Time** of instruction in SFT is short—from hours to a few weeks—vs. many weeks for a course in FHE. The **Format** of SFT is typically focused on some form of active learning vs. more traditionally lecture-heavy FHE. **Prerequisites** are easier to define and enforce in FHE, unlike in SFT. **Learner preparation** is also difficult to predict in SFT, vs. FHE where learners will have been predictably prepared by prior coursework or the course is designed to be foundational. **Instructors** in SFT are often domain experts but may have limited teaching experience; FHE instructors will usually have some expectation of preparation for teaching, may have been teaching the material regularly, and often have the benefit of access or being expected to use expert assistance in the planning and delivery of instruction. **Regulations**, policies, and laws usually apply to FHE courses; frequently, the informal nature of SFT is not affected by these and in practice SFT may not be regulated the same way as formal classroom instruction. **Sequence** of classes in a FHE curricular program provides learners with clear guidance on next steps, whereas SFT learners must direct their own learning; there may be additional SFT opportunities that can assist them in achieving their objectives, but this is not common. **Economics** of FHE assigns a concrete value to instruction; features of instruction, such as quality, can command a higher price; incentivizing maximized quality. Instructional effort is generally compensated. SFT instruction may be made available without cost to learners but this may result in an underestimation of the value or quality of instruction; there may be no economic incentives for optimizing “free” instruction. Instruction delivery may rely on uncompensated volunteer labor. **Variability** is the summation of all these characteristics. The variability of SFT is high (18), two courses on a similar topic may differ widely; instructional practices and curricula may not be documented for reuse. In FHE, variability in these characteristics is much lower; comparisons can be made across programs, allowing equivalencies and credit exchanges across institutions and programs. It is possible to make formal comparisons between distinct FHE programs (e.g., (34)) within one country or university system. Transferring from one higher education institution to another involves a systematic assessment of equivalence of prior work (e.g., (35); see also (36)). This figure emphasizes that strategies for improving FHE STEMM education would likely not be transferable to SFT without significant modification and that formal knowledge about teaching and learning is perhaps the only shared characteristic of these instructional forms.

Compared to FHE, the variability of SFT makes it far more difficult to address as a system. As noted by UNESCO (18), this variability arises on both the instruction side (“diversity, plurality, … decentralized and deregulated nature”) as well as from the learner side (*viz*. “variety of learners’ aims”). Except for a few large-scale SFT programs with an explicit focus on instructional quality, SFT instructors may have little knowledge or understanding of learner preparedness and contexts. SFT instructors are generally chosen for domain expertise and may not have pedagogical training or support that could help them adapt their teaching to overcome obstacles they encounter. Additionally, SFT courses are often bespoke, independent, and transient. This increases the chance that even effective instructional content and practices are unimplemented, unshared, and difficult to replicate. For learners, it may be impossible to compare different SFT opportunities on the same topic, meaning that decisions to enroll are based primarily on what is available. Learners wishing to prioritize effectiveness, accessibility, and inclusivity of instruction may also lack information or assurances on these characteristics in advance of the training. Overall, SFT lacks the stabilizing pedagogical, programmatic, policy, and economic structures that make improvement in the FHE context more tractable. Any given SFT course is therefore at risk of being a “black box,” having a definite form (i.e., short) but unknown contents (e.g., effectiveness and inclusiveness).

Since SFT lacks the system-context that is crucial to FHE reform (e.g., Reinholz et al. (30)), SFT reform could benefit from an approach that systematizes SFT. Rather than imposing FHE structures on a vastly different instructional context, systematization could be achieved by identifying features SFT programs have in common and designing interventions that address problems from multiple angles. For SFT, it is reasonable to conclude that reform efforts should engage the entire set of stakeholders (e.g., learners, instructors, instruction designers, administrators, funders) that make up the SFT “system.” Reforms that are actionable for both individuals and collectives have more possibilities for implementation. Recommended changes could be designed as standalone measures (e.g., a change an individual instructor could implement), or achieve impact as groups of people adopt (e.g., shared sets of standards or credentials).

Optimizing SFT for effectiveness and inclusion across the career span is timely and justified. The U.S. National Science Foundation 2026 Idea Machine project (NSF 2026) identified “high impact grand challenges” in research and STEM education that could help “set the U.S. agenda for fundamental research (37).” The research presented here emerged from the “Reinventing Scientific Talent” proposal, which was selected as an NSF 2026 grand challenge. This proposal called for the “transform[ation] of the education of scientists and STEM professionals after their formal training.” A small think tank-style conference sponsored by NSF 2026 (DRL#2027025) assembled representative global efforts in SFT to generate actionable recommendations for improvement.

## RESULTS

A two-stage conference assembled recognized experts in SFT, FHE, and the cognitive and educational psychology of higher education. Participants included several of the largest SFT programs reaching life scientists today (e.g., *The Carpentries* (38), *ELIXIR* training (39), *Galaxy Training Network* (40), *Cold Spring Harbor Laboratory* and *DNA Learning Center* courses (41, 42)), as well as emerging programs in SFT training, related research, and funding. Participants’ experience included SFT program development, deployment, and revision, as well as expertise in disability, equity, and inclusion. Some participants are engaged within their home countries, but most work overseas training in international settings. The content areas in which the participants’ training efforts are focused include bioinformatics, computational biology, computer science, data analysis, data science, genetics, molecular biology, and STEMM education in undergraduate and/or graduate contexts. There were 30 participants from ten countries: U.S. and Puerto Rico, Australia, Canada, France, Germany, The Netherlands, New Zealand, Sweden, Switzerland, and the United Kingdom, with similar participation for the in-person conference. Participant-reported demographic information included Gender: 73% Women, 23% Men, 3% Non-binary; Race/ethnicity: 73% White, 10% Asian, 10% Black, 3% Hispanic, 3% Indigenous; Other categories: 23% Underrepresented in the sciences, 7% disabled, 7% other identities. Post-conference, participants detailed the reach of their respective SFT programs. Annually on average, participants’ programs train 342 estimated new trainers (range: 0-7000), and train or reach an average of 1283 estimated learners (range: 0-8000). Among participants, *The Carpentries* SFT program reaches the largest number of countries: 64 (range: 1-64)(43).

This group derived 14 actionable recommendations (Tables 2–5) for systematic, evidence-based improvements to SFT aligned with the Bicycle Principles framework (see Methods). The Bicycle Principles (Figure 2) were developed by the project principal investigators with discussion among the organizing committee. Through discussion and minor modifications for clarity, it was determined by participants that The Principles accommodated all recommendations. The outputs from the virtual kick-off sessions converged on three areas:

1. **Catalytic Learning**: How SFT can better position learners to be self-directed after the completion of a training event (44).
2. **Inclusion**: How SFT can be made more inclusive for learners of diverse backgrounds and abilities.
3. **Scaling and Sustaining**: How, and with the help of what incentives, effective SFT can be sustainably scaled to large numbers of learners by large numbers of instructors.

**Table 1:**
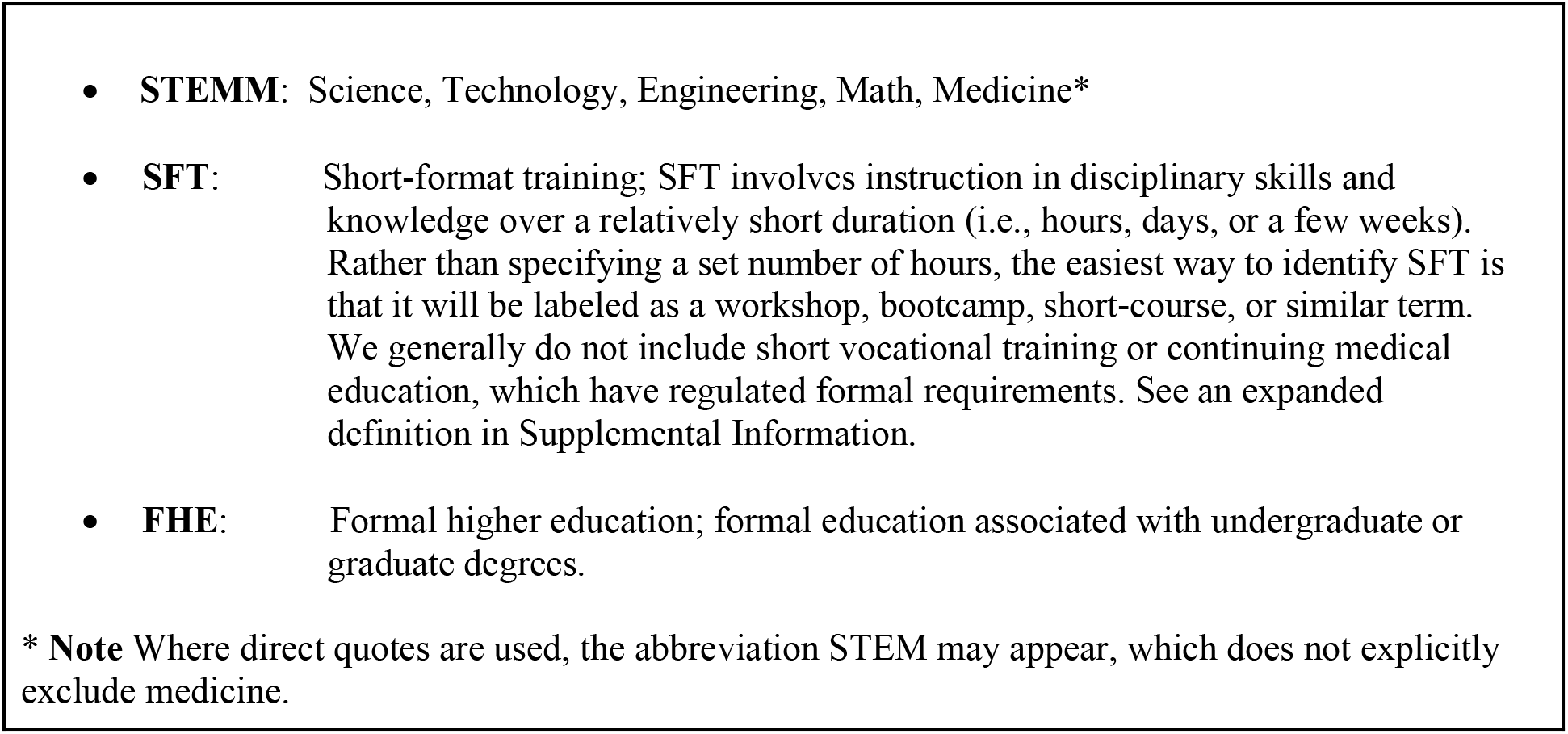
Abbreviations and definitions.

**Table 2.**
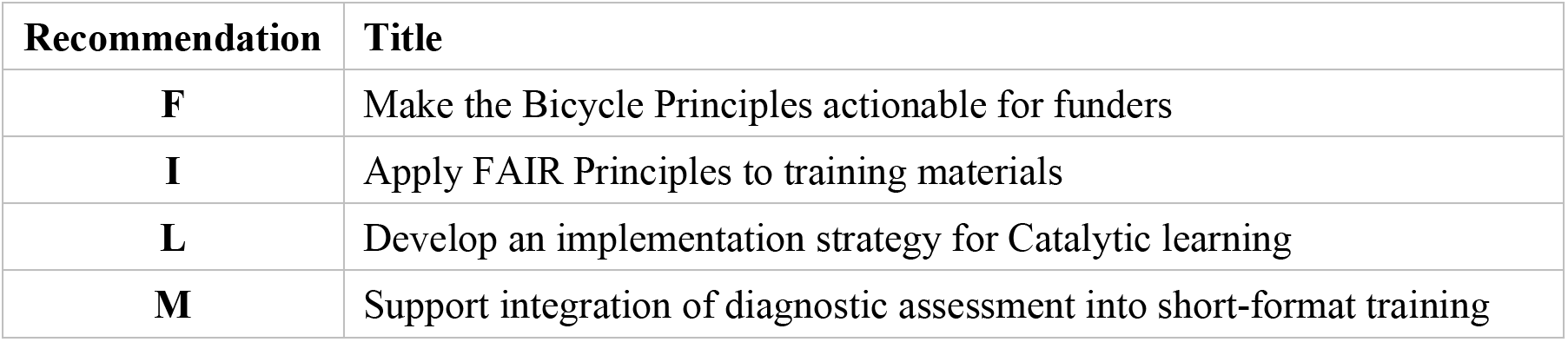
Recommendation Starting Points: Individual SFT Instructors—Grass Roots. To be actionable at the community or organization levels, these recommendations (I, M, L) include actions that individual instructors can take. Data from instructor’s own courses or scholarship could help deliver empirical results that justify future broader recommendation implementations. These data are likely needed to convince funders so recommendation (F) may also start with individual actions.

| Recommendation | Title |
| --- | --- |
| **F** | Make the Bicycle Principles actionable for funders |
| **I** | Apply FAIR Principles to training materials |
| **L** | Develop an implementation strategy for Catalytic learning |
| **M** | Support integration of diagnostic assessment into short-format training |

**Table 3.**
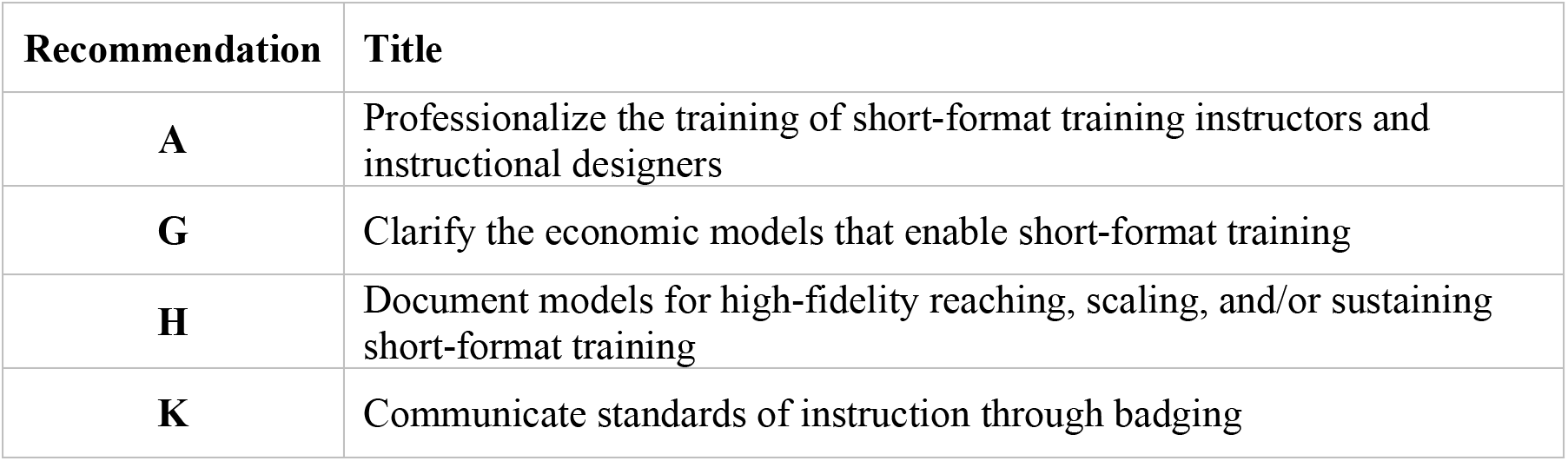
Recommendation Starting Points: Group/Community—leading from the middle: Community or group efforts (i.e., groups with primarily volunteer, rather than contractual, arrangements with members) are needed for recommendations (A, G, H, K) because without community endorsement, individual practitioners might find it difficult to achieve the broad buy-in needed to rally SFT instructors around shared goals and standards. Moreover, groups and communities currently engaged in professionalizing the training SFT instructors (e.g., *The Carpentries*, *ELIXIR*) might be best positioned to create and communicate standards of instruction through badging as well as describe economic models that underpin their SFT programs.

**Table 4.**
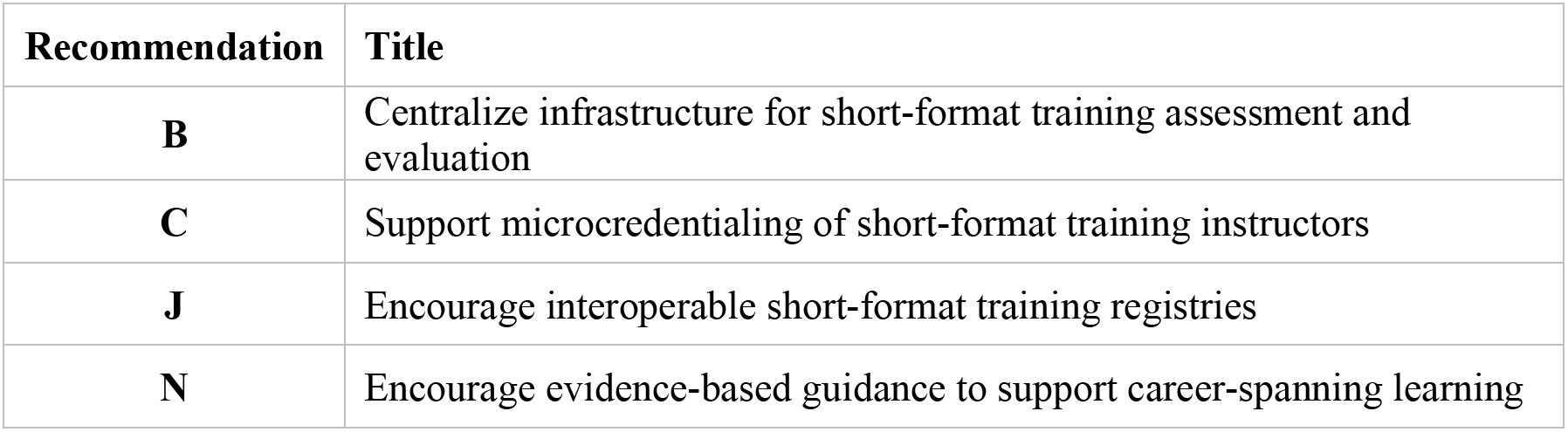
Recommendation Starting Points: Organizations and Institutions—Top-down: Organizations (e.g., employers or groups with formal contracts or understanding with individuals, groups/communities, or other organizations) would be able to implement complex recommendations which require teams of experts and sustained funding. These same groups can advance other recommendations, once grassroots efforts have laid groundwork.

**Table 5.**
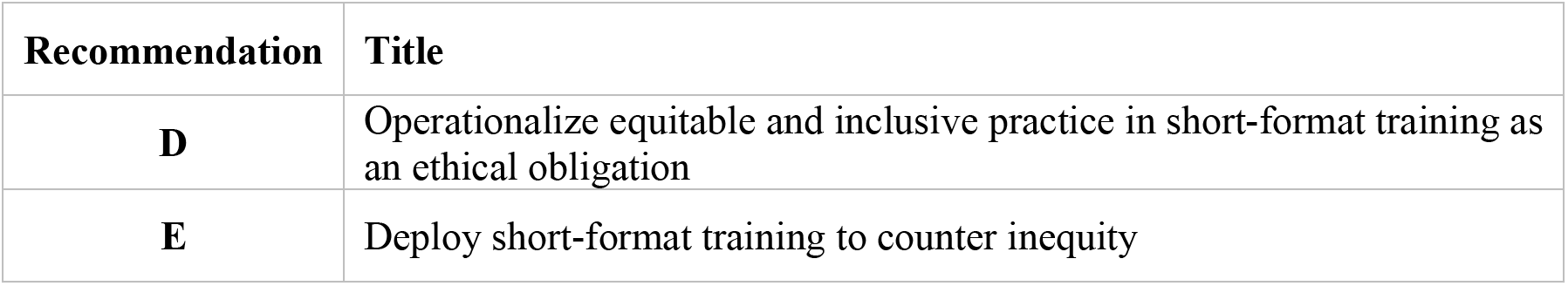
Recommendation Starting Points—Action at all levels: We identify that there are fruitful actions to be taken at all levels to implement these two recommendations.

**Figure 2.**
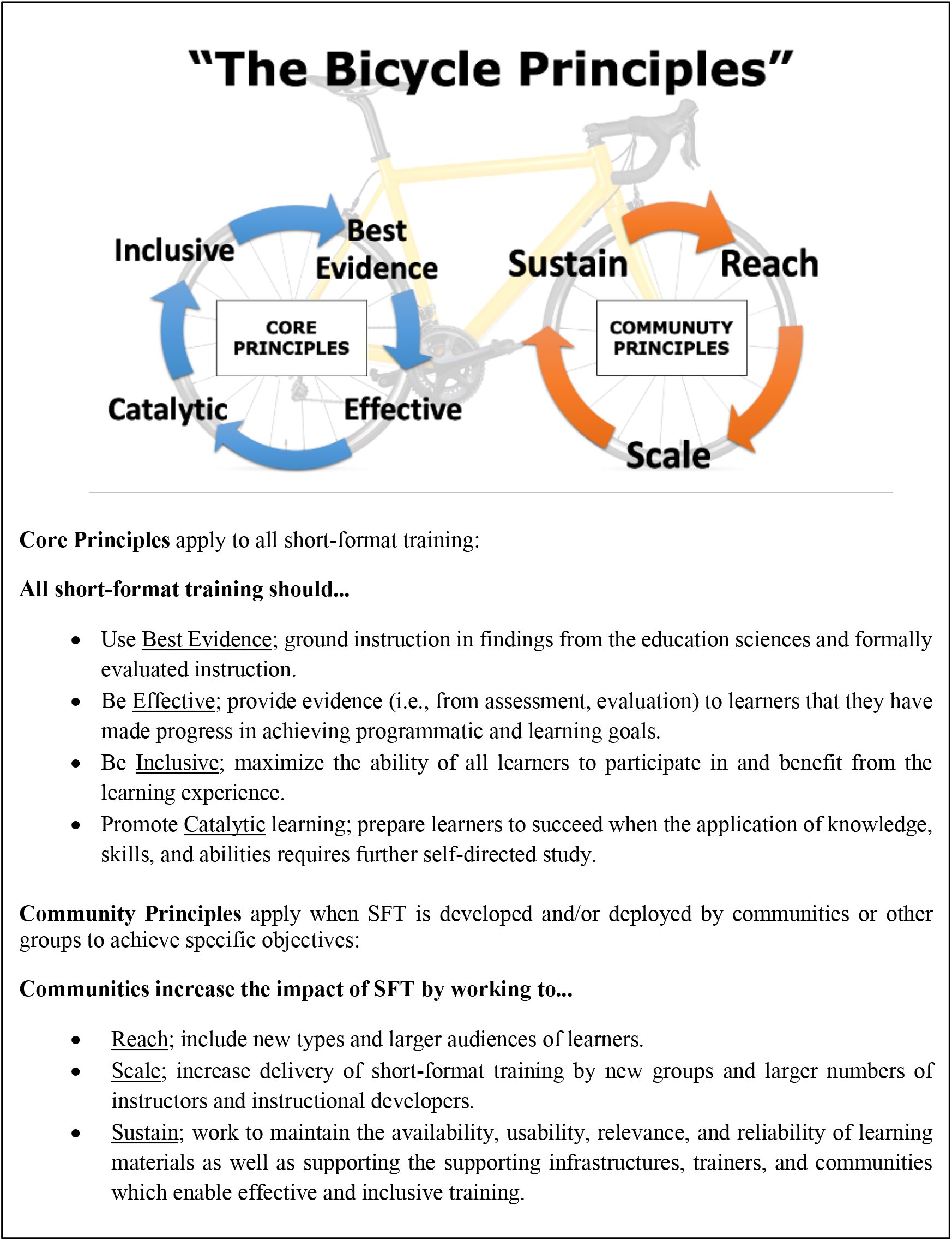
The Bicycle Principles: “The Principles” represent a framework for recommendations to improve SFT. One cycle comprises “Core Principles” that all SFT should meet: *Best Evidence*, *Effective*, *Inclusive*, *Catalytic*. The second cycle of “Community Principles” apply when the SFT is developed with the potential or intention to be reused and disseminated beyond the original or initial deployment: *Reach*, *Scale*, and *Sustain*. These two iterative cycles make up the “bicycle.”

At the hybrid conference, participants used a reproducible process (see Methods) to develop recommendations for improving the efficacy and inclusivity of SFT across the career span in alignment with the Bicycle Principles.

The group generated a document outlining 14 recommendations, where each recommendation is elaborated through six descriptive sections: 1) *Summary* expands upon the problem the recommendation tries to solve; 2) *How might this work* presents an implementation example and suggestions on evaluating success; 3) *Related Principles* lists closely related Bicycle Principles; 4) *Benefits to the learners* lists how recommendation helps learners (directly or indirectly); 5) *Incentives to Implementers* lists motivations for implementers to enact this recommendation; and 6) *Barriers to Implementation* lists obstacles that may hinder implementation of this recommendation. We provide a full example for Recommendation (A) in Supplemental Information and the entire set of recommendation descriptions is available at bikeprinciples.org (45). A Roadmap document (46) was also created, together with suggestions for implementation of each recommendation. The 14 recommendations do not have an intrinsic ordering, and when considering which stakeholder types might have the greatest success with implementation, groupings emerged (i.e., SFT instructors, professional groups and communities of practice, formal organizations and institutions, all stakeholders). The recommendations organized by suggested initial implementation stakeholder are summarized in Tables 2–5. See Supplemental Information for an alphabetized list of the recommendations.

## DISCUSSION

This work addresses unanswered calls to improve SFT efficacy (9, 10, 15, 18, 37) as well as the pressing need for educational reforms to improve equity and inclusion (11, 18, 47). We also underscore the increasing need to extend the attention of STEMM reform beyond undergraduate and graduate education. The NSF 2022-2026 strategic plan (48, p. 32) calls for “research that will develop and test new models for the lifetime integration of career and technical training, to keep pace with the ever-expanding frontiers of knowledge” and states that research on “how learning can continue throughout a person’s lifetime is crucial if we are to exploit these opportunities and maintain a competitive economy (p. 16).” The Bicycle Principles and 14 recommendations can support such research as they present testable assertions with evaluable impacts on learners.

Cognizant of the difficulties FHE STEMM reform has faced, and that SFT’s variability makes it even more complex to improve, there are two questions we should consider. First, why might this effort succeed in improving SFT in general? Second, what can be done to increase the likelihood that the Bicycle Principles and the recommendations are used?

The Bicycle Principles and recommendations can succeed because they serve as tools for making SFT measurable and standardizable. Requiring metrics and standards are a necessary step for any reform, and establishing a common set of principles creates reference points without insisting on rigid inflexibility. Without evaluable definitions for effective and inclusive SFT, it would be impossible to determine if and to what extent any change effort is successful. To quote an aphorism, “If you can’t measure it, you can’t improve it.”

The core Bicycle Principles demand that SFT is grounded in evidence-based teaching (*Best evidence*) and measured by evaluation and assessment (*Effective*). These Principles are explicit in several recommendations (e.g., B, C, H, K, M) and are consistent with curriculum and instructional guidelines (49).

Furthermore, The Principles require instructors to consider if SFT is an appropriate format for instruction. SFT should not be used—at least as the sole mechanism for instruction—when it is not compatible with the intended learning outcome(s). Incremental updates or “just-in-time” training is very compatible with SFT. Learners seeking complex sets of skills or retraining for proficiency in a new discipline have more complex needs and may need more than a “short” amount of time. The *Catalytic* principle and recommendation (L): *Develop an Implementation Strategy for Catalytic Learning,* suggests that instructors work to support self-directed learning when learning outcomes exceed what SFT can deliver on its own. Recommendations (H, I, J, M) would support learners in identifying additional SFT and other learning materials that could help them after an introductory training.

The final core Bicycle Principle (*Inclusion*) and the related accessibility and equity requirements must be prioritized since SFT’s short duration and less formal context often leaves these features neglected. Inclusion is meant to be a blanket concept that, ideally, applies to all persons or groups. Inclusion means creating an environment where everyone feels welcome, valued, respected, and has equal opportunity for equivalent participation. In practice—particularly in the context of STEMM research and training— creation of inclusive environments may fail to consider the needs of all groups (50). In situations where design is not co-developed, persons with disabilities are often left out of the inclusion conversation. Therefore, it is necessary to also define accessibility as “an umbrella term for all aspects which influence a person’s ability to function within an environment (51).” Accessibility is the design and implementation of systems, policies, processes, ways of interacting, and environments to ensure that persons with disabilities have equivalent access to a given space, and therefore equivalent experiences when participating in an activity. These observations underpin Recommendation (D): *Operationalize Equitable and Inclusive Practice in Short-format Training as an Ethical Obligation*. Implementation of this recommendation could inform and support instructors with tools that help them support equity, inclusion, and accessibility and develop the mindset that these features are a minimum standard for professional practice. We also know that pervasive inequities and disparities in FHE continue to harm STEMM professionals even after they overcome barriers to advanced degrees (52). Here, there is a positive opportunity to use SFT to correct disparity. Recommendation (E): *Deploy Short-format Training to Counter Inequity*, advocates for directing SFT resources to peoples who have been historically excluded from STEMM (e.g., minoritized ancestry groups, the disabled, low-income groups, the Global South). SFT resources should be thoughtfully and meaningfully deployed to counter inequity that may have resulted in historically excluded STEMM practitioners not receiving training opportunities that were available to others. In all these areas, actions are possible at all levels, and solutions must be co-created with the people they are intended to benefit.

A secondary reason why the Bicycle Principles and recommendations can succeed is that they work across the breadth of SFT, treating it as a system. If we consider the SFT “system” to be composed of its stakeholders, then we can impose some systematicity by developing stakeholder-focused solutions (e.g., Tables 2–5; see also Roadmap) since stakeholder groups are one of the few features all SFT shares.

Confidence that stakeholders will use The Bicycle Principles and recommendations relies partly on their origin from within the community, partly from their alignment with ongoing SFT activities worldwide, and partly from the structure they offer to those who seek to improve SFT. These factors make change plausible. Reinholz et al. (30) concluded that two change theories are the most commonly used in FHE reform: Community of Practice (53) and Diffusion of Innovation (54). Examples of SFT activities worldwide demonstrate that The Principles and recommendations are supportive of both theories. For example, *The Carpentries* SFT instructor training program (55) represents a community of practice consistent with The Principles and several recommendations (e.g., A, B, K, M). *The Carpentries* trainer curriculum requires instructors to be trained according to a set of evidence-based teaching standards, to integrate assessment into their two-day courses, and participate in discussion and feedback sessions to earn and maintain a credential. *ELIXIR-GOBLET* instructor training (56) and related *ELIXIR* training resources present examples of the Diffusion of Innovation theory, consistent with The Principles and recommendations (e.g., A, B, I, N); innovative instructional tools such as the Bioinformatics Mastery Rubric (57) provide guidance for career-spanning learning, various workshops and professional forums are opportunities for instructors to be exposed to knowledge about a new method, persuaded by its benefits, and supported to implement, customize, and adopt. Using the Bicycle Principles as a framework to improve SFT creates the opportunity to learn from FHE reforms—making what could work within the structured FHE environment more transferable to SFT.

Change theories in FHE reforms differentiate between changes that come about from top-down policies or emerge from individual or group actions (31). The Principles orient all stakeholders to a common set of objectives, such that the recommendations can be partitioned into individual, collective, and policy-based actions (i.e., Tables 2–5). Several recommendations could result in policies or strategies that are prescribed top-down (e.g., recommendations: B, C, G, K, N), but many recommendations achieve their greatest impact through wide-spread adoption by individuals (e.g., A, E, H, I, L). We appreciate that the recommendations assume a level of autonomy and community engagement that might not be plausible for every potential implementer—most recommendations cannot be implemented by individual SFT instructors alone. Just as in FHE reform, SFT instructors have responsibilities to enact some changes (e.g., Recommendation: D), but success is unlikely if the burden of change rests exclusively with instructors (30). Future work, including updates and customizations to a proposed Implementation Roadmap (46) will require creating a variety of approaches to bring recommendations into practice (e.g., checklists, instructor training, supportive infrastructures, policy mandates).

Finally, we note that although this work represents a consensus of experts, consensus cannot capture every possible circumstance. We leveraged the global reach of our organizing committee to recruit self-nominations, but limitations including COVID-19 meant we lacked direct representation from individuals in Africa, South and Central America, or Asia. However, we did have representation from organizations that have membership and activities in these regions. Starting in July 2022, the Bicycle Principles and recommendations have been widely disseminated online through bikeprinciples.org and through international conferences. Given the reach of the assembled group, and that online, in-person, and asynchronous dissemination, as well as focus groups have not surfaced any new or unaccommodated concerns, we believe the consensus derived by our group is likely to represent saturation on the topics. We do not take this to mean that more recommendations are not possible, only that we have arrived at a coherent set of recommendations. Within FHE STEMM education improvement efforts, a consistent finding is that success for such initiatives depends on considering the entire system in which the instruction occurs. Although Biswas et al. (33) and Reinholz et al. (30) are discussing FHE and undergraduate STEM(M) improvement, the SFT subject matter experts at our meeting identified recommendations for SFT-specific improvement that are similar to FHE-based guidance, albeit without the system-level structure of FHE. This post-hoc triangulation strengthens confidence in the validity of the results of this conference, while also highlighting the challenges facing individuals and communities in improving SFT.

To increase the likelihood that the Bicycle Principles and any recommendations are used, action is required at all stakeholder levels (30). FHE reform efforts have engaged stakeholders in several ways including funded research programs and institution-wide improvement projects. Journals and professional societies support dissemination of improvements. These mechanisms support SFT to a lesser extent; currently there is no comparable research program dedicated to SFT. SFT also lacks incentives that encourage innovations to be published, or that reward and recognize SFT instructors’ accomplishments.

Despite fewer formal incentives, there is evidence that communities of practice could be a valuable mechanism for promoting adoption of the Bicycle Principles and recommendations. The “community” set of Bicycle Principles prompt SFT programs to think about how materials could be shared, and instructors recognized and incentivized. For example, *LifeSciTrainers* (58) is an informal online community of practice for individuals engaged in SFT in the life sciences. Through it, instructors meet monthly and use online forums to share ideas and materials independent of SFT instructors’ affiliation with a specific program or topic area. Talk series highlight instructors’ accomplishments and provide an opportunity to share innovations in an informal setting. *LifeSciTrainers* activities are consistent with advancing recommendations (A, C, G, H, I, J), and provide an example of approaches that could help share effective practices across programs. International participation in *LifeSciTrainers* suggests global enthusiasm for SFT communities.

Funders must also exercise their role in promoting SFT reform. Over time, and consistent with recommendation (F), grassroots efforts could provide the evidence that justifies funders in imposing topdown standards for the effectiveness and inclusiveness of the SFT they invest in. Recent successes can be emulated. The FAIR Principles (59) were proposed in 2016 to reform scientific data management, a highly complex and multidimensional topic (e.g., technology, policy, incentives). The FAIR Principles were widely adopted by stakeholders and were enshrined in institutional policies globally, including the U.S. National Institutes of Health (NIH) in 2023 (60).

Ultimately, the final and most important group to involve in SFT reform will be learners. Learners who are empowered to insist on quality would be a powerful force for change. Every learner should be able to expect effective and inclusive instruction. An important aim of the Bicycle Principles and recommendations is to transform SFT from a “black box”—a learning experience where learners are uncertain about efficacy and inclusion to “back of the box”—a learning experience where implementations of The Principles serve as standardized and informative consumer “labels” which offer interpretable information on the efficacy, inclusion, and quality of instruction. Standardized, easy-to-compare SFT, would also benefit instructors and SFT funders.

## CONCLUSION

SFT improvement is urgent and achievable. As Demming and Noray concluded “there is a widespread perception that STEM workers are in short supply … but it is the new STEM skills that are scarce, not the workers themselves (1).” The Bicycle Principles and associated recommendations organize what education research and the most effective SFT programs have learned, providing a rallying point for global SFT improvement efforts. SFT reform is a strategic long-term investment in the STEMM professionals we have spent decades developing, could accelerate the pace of discovery, and could broaden participation in STEMM. The rapid evolution of STEMM disciplines calls for optimizing SFT to make it more reliably effective, inclusive, and career-spanning.

## METHODS

The study was approved (exempted) by the Georgetown University Institutional Review Board (IRB# STUDY00003859); To structure conference discussions, project PIs (J.J.W., R.E.T.) synthesized a draft set of principles from literature and experience which were further refined by the Organizing Committee (Organizers: J.J.W., R.E.T., and B.B., S.S.D., K.J.L., T.M., T.K.T., C.vG.). The Bicycle Principles synthesize education science and community experience into a framework for improving SFT through two cyclic (hence “bi-cycle”) and iterative processes (see Figure 2).

### Recruitment

Participants were recruited through a widely advertised self-nomination process and by direct invitation. The announcement was distributed to colleagues, communities of interest, and through social media (see Supplemental Information). Nominations were accepted April 14^th^ through May 31^st^, 2021, using a form also completed by direct invitees (see Supplemental Information). Through literature search, PIs identified and contacted 31 additional candidates with relevant expertise. Excluding conflicts of interest, organizers scored and ranked applicants. Participants who increased non-overlapping areas of expertise and added to gender, ethnic, and racial diversity were prioritized. A small number of participants (virtual and in-person) from policymaking and funding agencies or bodies were also recruited. Participants also included representatives from large scientific professional societies as well as private sector companies. In areas lacking representation (e.g., ethnicity, experience), additional invitations were sent, and selection concluded by September 2021. Our budget supported 20 in-person participants and virtual participants up to the intended cap of 30-35 total (to encourage full participation in discussion). Notably, recruitment, nomination, and selection process occurred during the COVID-19 pandemic limiting potential participants.

### Meeting 1 (100% virtual)

Although we originally planned a single meeting, this virtual kick-off meeting took advantage of the postponement of the in-person conference due to COVID-19. The organizers generated 20 vignettes (see Supplemental Information) on challenges associated with SFT based on content analysis with phenomenography of the vignettes compiled from nomination forms, together with others synthesized from the experiences of the PIs (J.J.W., R.E.T.). Prior to the kick-off, participants provided feedback on the accessibility of the virtual meeting tools. Work was captured in virtual whiteboards and Slack chat. Participants also received a *precis* (see Supplemental Information), the vignettes, literature underpinning The Principles (including (44, 49)), and summarized conference goals. To accommodate most time zones, two sessions were held via Zoom in December 2021. During the kick-off participants selected vignette(s) aligned with their interests and expertise. Next, participants developed and justified recommendations they felt could enhance SFT’s effectiveness, inclusiveness, relevance across the career span, or some combination of these features. The PIs examined all kick-off meeting outputs and applied content analysis to discern emergent themes.

### Meeting 2 (70% in person, 30% virtual)

In May 2022, a three-day hybrid conference was held at the Cold Spring Harbor Laboratory Banbury Center in New York. Participants gave 14 presentations on how The Principles and specific recommendations had been or might be implemented within their various represented programs or presented feedback in the context of their professional areas of expertise. Synthesized challenge vignettes (see Supplemental Information) presented during the kickoff were used as a starting point to develop and refine recommendations. The conference included an active, two-phase, Delphi-style assessment of each recommendation by participants, where judgements were elicited together with evidence supporting implementation of the recommendation(s), e.g., experience of participants with similar recommendations, or from the literature. Additionally, details on what “success” might look like if the recommendation were implemented, as well as barriers, incentives, and other considerations for each recommendation across multiple stakeholder groups. Conference outputs included notes compiled by participants and a dedicated science writer (N.C.) (15,488 words, plus additional comments); three participant-synthesized papers on Catalytic learning, Inclusion, and Scaling/Sustaining training (A.L., G.S.M., L.P., M.S., S.D., and J.J.W., R.E.T; 13,760 words, plus additional comments). These were again subjected to content analysis with ongoing member check in, which generated a recommendation synthesis document (10,944 words, plus additional comments). Extensive documentation and participation by all helped ensure saturation with respect to recommendations on each vignette and established that recommendations were sufficiently detailed for actionability.

## ACKNOWLEDGMENTS

This material is based upon work supported by the National Science Foundation under DRL/EHR: 2027025. Any opinions, findings, and conclusions or recommendations expressed in this material are those of the author(s) and do not necessarily reflect the views of the National Science Foundation. The authors wish to thank Rebecca Leshan and the staff of the CSHL Banbury Center for their guidance and facilitation of our convenings, and other participants in the conference including Charla Lambert.

