## Supplemental Information for "Optimizing Short-format Training: an International Consensus on Effective, Inclusive, and Career-spanning Professional Development in the Life Sciences and Beyond"

##### S1. 14 Recommendations for optimizing SFT

The 14 recommendations do not have a single intrinsic ordering and may be grouped by stakeholder (as in the main paper), topically, or in other ways. See **S3** for a full description of recommendation (A) and www.bikeprinciples.org for a full description of all recommendations and expanded definitions.

A. Professionalize the training of short-format training instructors and instructional designers.

B. Centralize infrastructure for short-format training assessment and evaluation.

C. Support microcredentialing of short-format training instructors.

D. Operationalize equitable and inclusive practice in short-format training as an ethical obligation.

E. Deploy short-format training to counter inequity.

F. Make the Bicycle Principles actionable for funders.

G. Clarify the economic models that enable short-format training.

H. Document models for high-fidelity reaching, scaling, and/or sustaining of short-format training.

I. Apply FAIR principles to training materials.

J. Encourage interoperable short-format training registries.

K. Communicate standards of instruction through badging.

L. Develop an implementation strategy for Catalytic learning.

M. Support integration of diagnostic assessment into short-format training.

N. Encourage evidence-based guidance to support career-spanning learning.

#### S2. Short-Format Training (SFT) expanded definition

This was the full definition of SFT developed and used by participants at the workshop.

Short-format training involves instruction in disciplinary skills and knowledge over a relatively short duration (i.e., hours, days, or a few weeks). Rather than specifying a set number of hours, the easiest way to identify SFT is that it will be labeled as a workshop, bootcamp, short-course, or similar term. SFT generally has the following features:

- Generally happens outside of a formal undergraduate, graduate, or other degree-granting program. Generally, SFT learners do not receive a summative grade or university course credits. SFT may also include post-degree training programs for professional accreditation (e.g., physicians or medical specialists training for fellowship exams in a specialty). Learners may be recognized as having earned “professional development” or “continuing education” credit.
- Content is determined by instructors or instructional designers, not necessarily by an institutional or professional society curriculum approval process (e.g., accreditation).
- Tends to be stand-alone, not requiring formal prerequisites or required subsequent SFT. There may be expectations for learner preparation or recommended prerequisite knowledge. SFT can be linked and can have prerequisites, but these characteristics are not enforced as they might be in a formal setting.
- Typically delivered to a group of learners who enroll because of their interest in the topic, rather than a mandate.
- Typically developed and delivered by domain experts outside of and separately from an institutional undergraduate/graduate coursework teaching role (if they have such a role).

The extent to which a specific SFT course meets any of the above features will vary.

#### S3. Complete Example: Recommendation (A) full description

Each of the 14 recommendations has several descriptive features:

- **Summary**: Expands upon the problem(s) the recommendation tries to solve.
- **How might this work**: Brief implementation example and suggestions on evaluating success.
- **Related Principles**: Most closely related Bicycle Principles.
- **Benefits to the learners**: How recommendation helps learners (directly or indirectly).
- **Incentives to Implementers**: Motivations for implementers to enact this recommendation.
- **Barriers to Implementation**: Obstacles that may hinder this recommendation.

We provide here an example of one full recommendation as developed by conference participants. The complete list of recommendations and descriptions are available at www.bikeprinciples.org. Please note this website also contains additional detailed definitions of terms
(see: https://www.bikeprinciples.org/#definitions/)

##### A. Professionalize the training of short-format training instructors and instructional designers

**Summary:**

Creating a professional society(ies) or communities of practice dedicated to SFT could support instructors, instructional designers, and instructional administrators, as well as help to extend the reach of the *Bicycle Principles*.

Instructors, instructional designers, and instructional supporters/administrators could benefit from services provided by professional societies (e.g., a venue for self-identification, shared affiliation; dissemination of work; development of practice; rewarding of outstanding contributions). Instead of being a "special interest group" splintered from a specific professional society, this could be an independent, cross-cutting, professional community that can promote appreciation for the professional competency of effective SFT instructors. It could also positively influence the entire scientific community by promoting effective and accessible training principles worldwide. This group could focus on training to assist SFT instructors and instructional designers to reach specific standards. Since most professional associations already have instruction-focused special interest groups, there is evidence many people would be interested. Furthermore, existing but unconnected groups around the world already have formal, iteratively-improving trainer-training programs in which new instructors engage.

**How might this work:**

A SFT training professional society could be jointly established by organizations and/or projects that have significant SFT interest, expertise, and activity. As a community of practice, the society could: 1) vet, curate, and/or maintain centralized resources; 2) work to clarify and address significant challenges; and 3) share innovations for the delivery of effective, inclusive, and career-spanning SFT. Like most professional societies there could be virtual and in-person convenings, topical or geographical sections, and various communication mechanisms. Participation in this community would support recognition of peers and the strengthening and promotion of SFT teaching competencies.

Success for this new professional society would look like a growing membership base and the co-production of actionable artifacts for others to reuse (e.g., recommendations, guidelines, methods). The unique mission of this society is its emphasis on improving the consistency and quality of SFT training, at the level of the individual instructor and instructional designers as a profession. Individuals from diverse scientific disciplines could join this society to signal their commitment to effective, inclusive SFT instruction.

**Related Principles:**

- All

**Benefits to the learners:**

1. Increases the likelihood of successful SFT learning experiences when instructors have benefited from training and knowledge exchange promoted by the SFT professional society.
2. More high-quality instructors trained by the society increases the chance of meeting learner demand for SFT.

**Incentives to implementers:**

*For Instructors*

1. Recognition of their role and its importance in the scientific community, peer support, and increased sense of community. Professionalization may counter the perception that SFT (which is often delivered at no cost to learners) is not of "real value".

*For Instructors, Instructional Designers, Instructional Supporters*

2. Collective impact could disseminate effective practices and enhance the adoption of practices that individual instructors or SFT programs might have difficulty implementing alone.
3. Opportunities to develop new, better, and sustainable economic models for SFT as well as models for applying community *Bicycle Principles* (i.e., *Reach*, *Scale*, *Sustain*).

*For Funders and Organizations*

4. Working with an SFT professional society may strengthen justifications for SFT funding and allow funders to interact with multiple programs through a single group. The professionalization of instructors brings value to SFT provided by organizations.

**Barriers to implementation:**

1. There must be sufficient global grassroots support to form such an organization, and this requires clear value propositions for members and financial sponsors.
2. There are legal, logistical, and financial barriers to launching an entirely new organization. It may be possible for existing organizations and nonprofits to help builders leverage existing resources and groups.
3. To be successful, this organization will have to demonstrate significant and positive effects on the community of instructors and instruction developers. Many of these community members offer SFT as simply a transient or minor activity. If the new organization can show benefits for them, it will strengthen the effort.

**S4. Draft challenge vignette list for kick-off meeting.**

Based upon our original proposal, consultation with the organizing committee, and discussion with community members, we identified an initial problem set that could potentially be addressed at the conference and distributed this list to all participants ahead of the kick-off meeting (meeting 1):

1. How can an instructor ensure that everyone who needs to know, knows about a learning opportunity?
2. People expect to be given high quality training for free, and simultaneously consider free to mean low quality; how can instructors resolve this conflicting "truth" about online training?
3. How can I integrate ethical content into my course?
4. How do instructors establish, gauge, and maintain learners' engagement?
5. How do instructors deliver content when the subject matter is in flux?
6. How do instructors begin developing a course when there don't seem to be pre-existing training materials on a topic?
7. How should SFT be structured so that learners who misinterpret or fail to meet prerequisites still have a good chance of a successful learning experience?
8. Can learners be helped to differentiate "good" from "bad" instruction before their SFT experience?
9. How can instructors know if their assessments are working?
10. How can instructors assess catalysis? Is it enough to add it, or do you have to assess it?
11. How could instructors advise learners with highly individualized needs about future learning? How important is it that instructors identify individuals with unique needs?
12. How might instructors know if their learners are succeeding *after* an SFT event?
13. How can learners determine if an SFT is appropriate for them?
14. How should instructors plan for diverse needs of learners?
15. How should instructors manage the learning environment to maximize participation, and identify and minimize barriers to engagement?
16. How can we facilitate learners to define their specific needs in SFT?
17. What should instructors do if they don't have time to design, deliver, and/or grade assessments?
18. What would make training materials more reusable for instructors? At what point in development should this be considered?
19. What are incentives to adopt/disincentives from adopting specific (evidence-based) teaching practices?
20. How could instructors increase the diversity of attendees who participate in training?

#### S5. Conference nomination form and community distribution

##### Nomination Form Questions

The following form was used to evaluate applicants to the conference:

### PREVIEW: SELF-NOMINATION FORM

For preview purposes only; only nominations submitted online will be considered.
Self-nomination form: URL

#### MAKING CAREER-SPANNING LEARNING IN THE LIFE SCIENCES INCLUSIVE AND EFFECTIVE FOR ALL

December 2021*, Banbury Center, New York, USA

[*By July 2021, those selected will receive official invitations including details about the meeting date/time, accommodations, and travel]

We are planning for an in-person conference; those selected for the meeting will receive an official invitation from the Banbury Center by July 2021. This invitation will include conference dates as well as full travel and accommodation details. We recognize that decisions may be contingent upon evolving pandemic-related circumstances.

The meeting will span four days/three nights: beginning at 6.00pm EST on Day 1, and concluding around 2.00pm EST on Day 4. Lodging and meals will be provided for all invited participants, and the Center will work directly with invitees on any non-standard accommodation/conference facility requests. A travel allowance will be provided for participants from academic and other not-for-profit organizations; this allowance typically covers all associated travel expenses.

While the majority of participants will be in-person, a limited number of VIRTUAL attendees will be accommodated. Though we are making virtual participation possible, we do not intend for it to be a substitute for in-person participation. At the time of invitations, the Banbury Center will work with each virtual attendee to ensure, on a case-by-case basis, that meaningful participation is possible. [Note: The meeting will be held in the US Eastern time zone].

Participants from policy-making and funding organizations may apply to attend as observers.

We expect all in-person attendees to be present for the entire duration of the meeting. Virtual attendees (and observers) will not be required to attend sessions during hours considered unsocial in your time zone, nor informal discussion periods.

This self-nomination form includes three sections:

I. Career & Education (selection or short answer)
II. Related Experience (short answer - 300 words maximum)
III. Demographic Information (selection)

You will also be asked to upload your CV (two-pages maximum) –OR—a statement of related professional experience (two-pages maximum). The CV or statement should be a PDF, and the file named using the format: LASTNAME_FIRSTNAME

For section II (Related Experience), we strongly recommend composing your answers in a separate document to track character count, then pasting the response into the field.

**The deadline for submission is MAY 31, 2021**

First Name / Given Name

Last Name / Surname

Email address [*This address will be used to contact you if you are selected for an invitation AND this address will receive a copy of your application upon submitting. Please 'whitelist' our address to ensure you are able to receive our messages.]*

If invited, how do you intend to participate?

- I intend to participate IN-PERSON
- I intend to participate VIRTUALLY
- I intend to participate as an IN-PERSON OBSERVER (e.g., representing a funding or policy-making body)
- I intend to participate as a VIRTUAL OBSERVER (e.g., representing a funding or policy-making body)

#### I. Career & Education

Current employer

Current position (job title)

Career stage

- Early-career (typically within 10 years of terminal degree)
- Mid-career (more than 10 years after terminal degree)
- Late-career or beyond (within 5 years of retirement, emeritus, or retired status)

Career status

- Non-tenure track
- Tenure track
- Tenured
- Non-academic
- Other (please specify)

Education: Highest degree(s) obtained

Please list up to five products of your work/experience relevant to the conference (e.g., publications, courses taught)

CV or statement upload: Two pages maximum (any text beyond a second page will be deleted); PDF format only; File should be named using the following format: LASTNAME_FIRSTNAME

#### II. Relevant Experience

How does your professional work relate to the topic of short-format training? (1800 characters maximum, approximately 300 words)

Are there additional areas of expertise or personal experience relevant to short-format training that you would bring to this conference?
If nothing additional, please enter "NA." (1800 characters max., approx. 300 words)

What community(-ies) would you represent at this meeting, and how would you bring the work done at the meeting back to a community(-ies)? (1800 characters max., approx. 300 words)

Why do you want to participate in this conference? What would this conference have to achieve to make it worth your time? How would you contribute to achieving this? (1800 characters max., approx. 300 words)

#### III. Demographic Information

The Banbury Center and conference organizers are committed to the values of diversity, equity, and inclusion. We believe this meeting's effectiveness will rely on convening a group of participants that reflects the diversity of those we expect our work/outputs to serve. Therefore, demographic information will be one of the factors used in selecting participants.

Please indicate your gender

- Woman
- Man
- Nonbinary
- Prefer not to disclose
- Prefer to self-describe (please specify)

Which category(-ies) describe your race/ethnicity?
*Select all boxes that apply; you may select more than one category.*

- White (e.g., German, Irish, English, Italian, Polish, French)
- Hispanic, Latino, or Spanish origin (e.g., Mexican or Mexican American, Puerto Rican, Cuban, Salvadoran, Dominican, Colombian)
- Black (e.g., African American, Jamaican, Haitian, Nigerian, Ethiopian, Somalian)
- Asian (e.g., Chinese, Filipino, Asian Indian, Vietnamese, Korean, Japanese)
- Indigenous, First Nations, American Indian, or Alaska Native (e.g., Aboriginal, Navajo Nation, Blackfeet tribe, Mayan, Nahua, Native Village or Barrow Inupiat Traditional Government, Nome Eskimo Community)
- Middle Eastern or North African (e.g., Lebanese, Iranian, Egyptian, Syrian, Moroccan, Algerian)
- Native Hawaiian or other Pacific Islander (e.g. Samoan, Chamorro, Tongan, Fijian)
- I prefer not to say
- Other (please specify)

In what country are you primarily employed?

Do you consider yourself any of the following?
(you may select more than one)

- A race/ethnicity underrepresented in the life sciences (e.g., Indigenous, Black, person of color)?
- A disabled person
- Another category with which you wish to identify
- Yes, but prefer not to specify
- Prefer not to respond
- Unsure
- Other (please specify)

If “Another category with which you wish to identify,” please specify

— END OF PREVIEW —

**Communities and Mailing Lists**

The conference and nomination process was circulated to the following groups and lists: The Carpentries (>3,000 volunteer instructors), CyVerse mailing list (>80,000 subscribers), ELIXIR, Australian BioCommons (>1,500 subscribers), and Galaxy Training Networks; NIH intramural training and extramural research offices, LifeSciTrainers.org blog, Open Life Science; the Global Organisation for Bioinformatics Learning, Education and Training (GOBLET); BioQUEST Curriculum Consortium (20,000 users and >6,000 subscribers); through the Co-PIs of the NSF RCN-UBE: Establishing a Genomics Education Alliance: Steps toward Sustainability (GEA) and affiliated networks like the Genomics Educational Partnership (GEP).

**S6. *Precis* (abbreviated) for kick-off meeting**

This summary was presented to conference attendees and contains additional background materials. Note that some terminology in this document evolved during the conference as reflected in the main paper.

**A Set of Principles for Making Career-spanning Learning in the Life Sciences Inclusive and Effective for All**

The rapidly-increasing interdisciplinarity of the life sciences makes career-spanning learning critical. Scientists who are able to recognize and traverse skills gaps are better positioned to pursue impactful science and achieve their personal career goals. To bolster skills, life scientists often turn to short-format training (e.g., workshops, boot camps, and short courses) for professional development. While short-format training (SFT) provides point-of-need help, and is often low in time commitment and costs, this approach can be less successful than typically assumed.

We propose a set of principles for ensuring that short-format training in the life sciences is effective and accessible to all. "*The Principles*" are intended to promote the professionalization of SFT and can support several aspirational outcomes:

1. Enable stakeholders to design (instructors), select (learners), or deploy (institutions, communities, and funders) effective, catalytic, and inclusive training opportunities in support of career-spanning learning.
2. Empower learners to reliably assess if a short-format professional development opportunity will help them achieve their learning goals by promoting the adoption of evidence-based practices from the learning sciences.
3. Establish inclusivity and "catalysis" (support for ongoing self-directed learning) as essential to short-format training.

Buy-in and application of these principles for SFT could influence how we approach career-spanning learning in the sciences, supporting our vision of career-spanning training that is inclusive and effective for all.

**The principles, challenges, and work of the conference**

We describe each of The Principles below with rationale and example questions that might arise in the design, planning, delivery, and refinement of SFT. We attempt to show that the principles we have identified accommodate a diverse set of challenges and ties to our purpose of advancing inclusive and effective career-spanning learning.

We also have listed potential challenges that we can address at the conference. The challenges (which include your suggestions and feedback) are meant to illustrate realistic problems SFT learners and instructors face. The set we've selected is open to expansion and refinement (see "evaluating the principles" at the conclusion of this document). During the December kickoff, we will focus on challenges that are directly within the purview of instructors to address. We also realize broader challenges exist; we should document them as we identify them and will have a separate approach for working on these at the May conference.

During kickoff, small groups will work on the challenges that resonate most with them; we do not anticipate addressing all possible challenges. An accompanying "worksheet" will present a structured process for each small group to refine that challenge, adapt the challenge (if needed) to a specific setting, and propose structured solutions that might address that challenge. The worksheet will prompt you to consider specific lines of evidence including drawing on the education research literature (see: Workshop Reading) as well as your own expertise and experience. Small-group discussions, evaluation against The Principles, reformulations of groups, etc. will hopefully lead to strategies and recommendations that will resonate with others.

#### CORE PRINCIPLES I-IV

The first set of principles (Best Evidence, Effective, Catalytic, and Inclusive) are referred to as "core" because we believe all SFT must address them. Numbering of the principles suggests a sequence (e.g., training needs to be effective before trying to maximize its inclusivity), but the temporal order is not necessarily strict, and all are required.

Designing SFT based upon best available evidence from the learning sciences (peer-reviewed literature, books, and experience) is the **first principle** since the creation of learning objectives is universally agreed to be the optimal starting place for effective teaching. Once lessons are created, evidence of their effectiveness—based on assessment that is aligned with the objectives and the instruction—is the **second principle**. Determining if SFT supports sustainable or ongoing, self-directed learning, which we have designated "catalysis", is a novel component to establishing effectiveness. Because SFT is typically intended to augment the life scientist-learner's skill set, and not just their foundational knowledge, we characterize instruction as effective if it both supports the learner's achievement of learning objectives but also supports ongoing learning and application of the new knowledge—i.e., if SFT is catalytic, then it serves its ostensible purpose. Thus, catalysis is the **third principle, expanding what we consider "effective" instruction beyond achieving the more immediately verifiable learning objectives**. We assert that training that cannot promote catalysis for learners is not fully effective. Inclusivity is the **fourth principle**, not because it is less important than others but because we believe disseminating ineffective training, however inclusive, is unhelpful and harmful. Some elements of inclusivity are applicable at the best evidence phase of instructional design, whereas different inclusivity features may be relevant for determining SFT effectiveness or catalysis. Finally, inclusivity requires ongoing refinements that may become apparent as more learners provide feedback over time.

Bicycle Principles:

**I. Best Evidence**
*SFT must be grounded in evidence-based practices from educational research.*

**II/III. Effective and Catalytic**

*SFT should be effective, providing evidence to learners that they have made progress in achieving learning goals. Learners should be assisted in formulating additional learning goals and prepared for future self-directed learning.*

#### COMMUNITY PRINCIPLES V-VII

The second set of features (Shareable, Scalable, and Sustainable) are called community principles because we posit that these emerge when groups and communities of individuals think about long-term impacts of training and the potential to build skills for large groups of people. Developers of SFT may have no intention of making their training broadly available by sharing (increasing the numbers of instructors) or scaling (increasing the number of learners). However, when training is shared/scaled these principles could be used to promote organized progress across the discipline.

##### V/VI/VII. Shareable, Scalable, and Sustainability with Fidelity

*Training that meets the Core Principles can be shared with new and larger audiences. This sharing should be done to maximize fidelity to the Core Principles even when adaptations are made.*

#### CONCLUSION

##### Evaluating the Principles

These Principles are a starting point for the conference and will hopefully provide a common point of focus for us to exchange ideas around. Are they complete? Can you think of any important principles that are missing or do not fit? Are these principles potentially a starting point for SFT instructors to coalesce around or buy in to? We must also consider the specific conditions of the life sciences and how these principles may apply in this and other scientific domains.
